# A bioluminescence resonance energy transfer (BRET) assay to detect telomere length in *S. cerevisiae*

**DOI:** 10.64898/2026.03.11.711003

**Authors:** Felix Richter, Honorata M. Ropiak, Jörg Urban, Jacqueline Franke

## Abstract

A method to measure telomere length in *S. cerevisiae* was developed based on bioluminescence resonance energy transfer (BRET). The system uses energy transfer between a luciferase-Rif2 fusion protein and fluorescently tagged Rap1. The study demonstrates that the BRET ratio correlates with the Rap1/Rif2 complex at the telomeres and thus the availability of telomeric Rap1 binding sites. This enables the measurement of telomere length in living cells. The system was able to reproduce reported deviations in telomere length in mutants lacking telomere length regulators, cells treated with telomere length modifying compounds and strains expressing inducible telomerase. The BRET ratio linearly correlated with the average number of telomeric nucleotides derived from long-read sequencing data using a novel algorithm for telomere length calculation.

**GRAPHICAL ABSTRACT:** 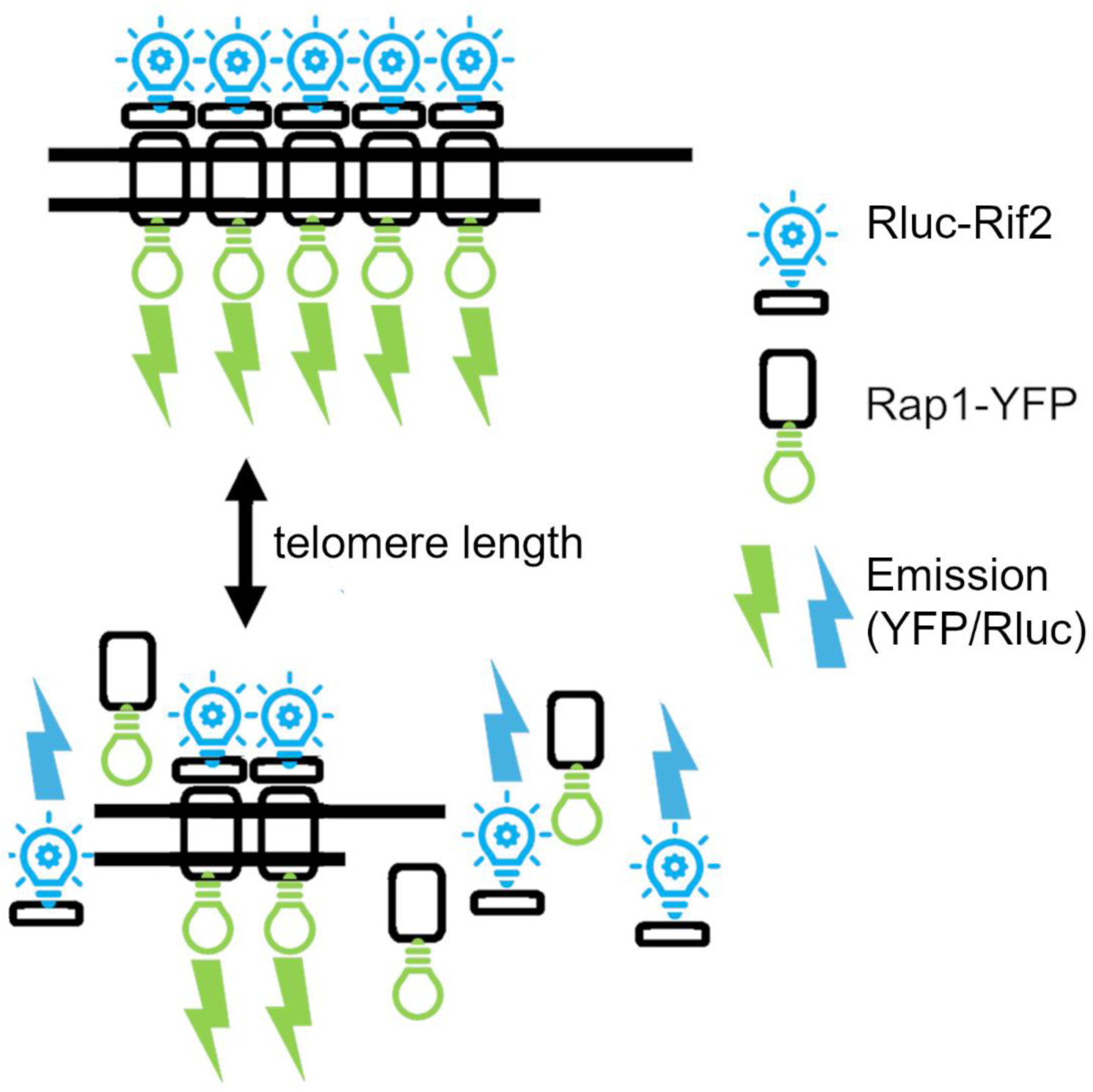

## INTRODUCTION

Telomeres are one of the guardian structures of genome stability. They are the terminal structures of chromosomes and are composed of double stranded telomeric repeats and a G-rich 3′-overhang^1^. They are occupied by several complexes of specific telomeric proteins that stabilize their three-dimensional structure, which loops back to the subtelomeric region^2^.

Telomeres avoid the recognition of the chromosomal ends as DNA breaks by the cellular repair machinery, circumvent degradation and recombination, and regulate the access of the telomerase complex which can elongate short telomeres^3^.

Most adult somatic cells in humans do not express telomerase and are subject to the end-replication problem, i.e. the loss of genetic material in each cell division due to end resection and the inability of the replication machinery to reach the 3′-end of linear DNA^4^. This limits their replicative capacity to the so-called Hayflick-limit at which at least one telomere is critically short. Cells that reach this limit go into permanent cell cycle arrest and cellular senescence^5^. This is one of the reasons for the accumulation of senescent cells in old age, contributing to reduced function and aging associated pathologies. For this reason, telomere attrition is one of the fundamental hallmarks of aging^6^.

An evolutionary advantage of the inactivation of telomerase in somatic tissues could be the role of telomeres as a firewall against cancer. When somatic cells acquire mutations or epigenetic alterations that let them expand clonally without normal inhibition, they soon reach their Hayflick-limit and go into apoptosis or senescence. Malignant transformation requires bypassing the Hayflick-limit. In a large majority of analyzed tumor biopsies telomerase activity could be detected^7^. Telomere maintenance as part of enabling replicative immortality is thus one of the hallmarks of cancer^8^.

It seems like a contradiction that disrupting telomere maintenance or shortening of telomeres is a possible target for cancer treatment while lengthening of telomeres is pursued as a possible geroprotective intervention, as shown in mice^9^. However, the main difference between cancer and functional telomeres is their length and stability which could make it safely possible to target very short cancer cell telomeres without greatly affecting the normal cell population and elongating telomeres in healthy aged individuals without increasing the cancer risk^10^.

Although compounds that increase or stabilize telomere length are potential means to increase health- and lifespan^11^ and compounds that shorten telomeres or inhibit their maintenance are promising cancer therapeutics^10^, the clinical success in both fields is still moderate. This is mostly due to severe side effects or low efficiency of the few candidate compounds^12^.

One reason for the low number of telomere affecting compounds is that so far there are no high-throughput capable methods to screen for changes in telomere length or dynamics. The classical methods to measure telomere length are terminal restriction fragment (TRF) analysis^13^, fluorescence in-situ hybridization (FISH)^14^ or quantitative PCR (qPCR)^15^. A recent method is based on long-read sequencing^16^. All these methods are time, labor and/or cost intensive. qPCR based methods can be used to measure a moderate number of samples, however for high-throughput screening the method is too elaborate and expensive and shows significant intra-and inter-assay variation^17^.

In summary, up until now, there was no high-throughput method to assess telomere length and dynamics.

The well characterized yeast *Saccharomyces cerevisiae* has been established as a powerful screening system as it provides a rapid workflow, high experimental feasibility, low costs and reproducible results. Although telomere length regulation is more complex in mammalian cells, it is known that certain mechanisms of telomere replication, regulation of telomere length and structure are largely evolutionarily conserved from yeast to mammals including various functional homologues^18^. Interestingly, studies in yeast have identified several small molecules that also show effects human cells^19^.

Here, we demonstrate a system that determines telomere length in *S. cerevisiae* based on protein-protein interaction using Bioluminescence Resonance Energy Transfer (BRET). The results show that the assay assesses telomere length quantitatively and independently from cell density. It is quick, cost effective, requires very little hands-on time and works in a 96-well-format with standard laboratory equipment. It is therefore a tool for basic research and high-throughput compound screening for new lead compounds to increase or decrease telomere length. These compounds could offer therapeutic benefits for cancer, aging associated pathologies and telomere biology disorders.

## RESULTS

Previous studies have shown that yeast Rap1 forms a more stable complex with Rif1 and Rif2 on arrays of more than four Rap1 binding motifs, that occur only on telomeric DNA^20^. Based on this, we hypothesized that, assuming constant levels of telomeric proteins, the ratio of Rap1 bound Rif2 to unbound Rif2 would be proportional to the number of available Rap1 binding sites. Consequently, the ratio of Rif2-Rap1 complex to unbound Rif2 would be proportional to the overall length of telomeric repeats in the cell.

To test this hypothesis, we genomically fused *RAP1* with the nucleotide sequence for Yellow Fluorescent Protein (YFP) and *RIF2* with the sequence for *Renilla reniformis* luciferase (Rluc) in *S. cerevisiae*. This tagging strategy enabled approximation of interaction between Rap1 and Rif2 by determining the BRET ratio, defined as the Rap1-YFP emission divided by the emission of Rluc-Rif2 (see Equation 1).

We then measured the BRET ratio in the tagged wild type *S. cerevisiae* strain and tagged deletion mutants known to possess shorter (*Δtel1*) and longer (*Δelg1*) telomeres^21^.

Fig 1A shows, the average BRET ratio in *Δtel1* cells was significantly lower than in wild type cells (1.11 ± 0.03 vs. 1.66 ± 0.05, 95 % confidence interval - CI) and the BRET ratio measured in *Δelg1 cells was* significantly higher than in wild type cells (1.86 ± 0.11 vs. 1.66 ± 0.05, 95 % CI).

**Fig 1.**
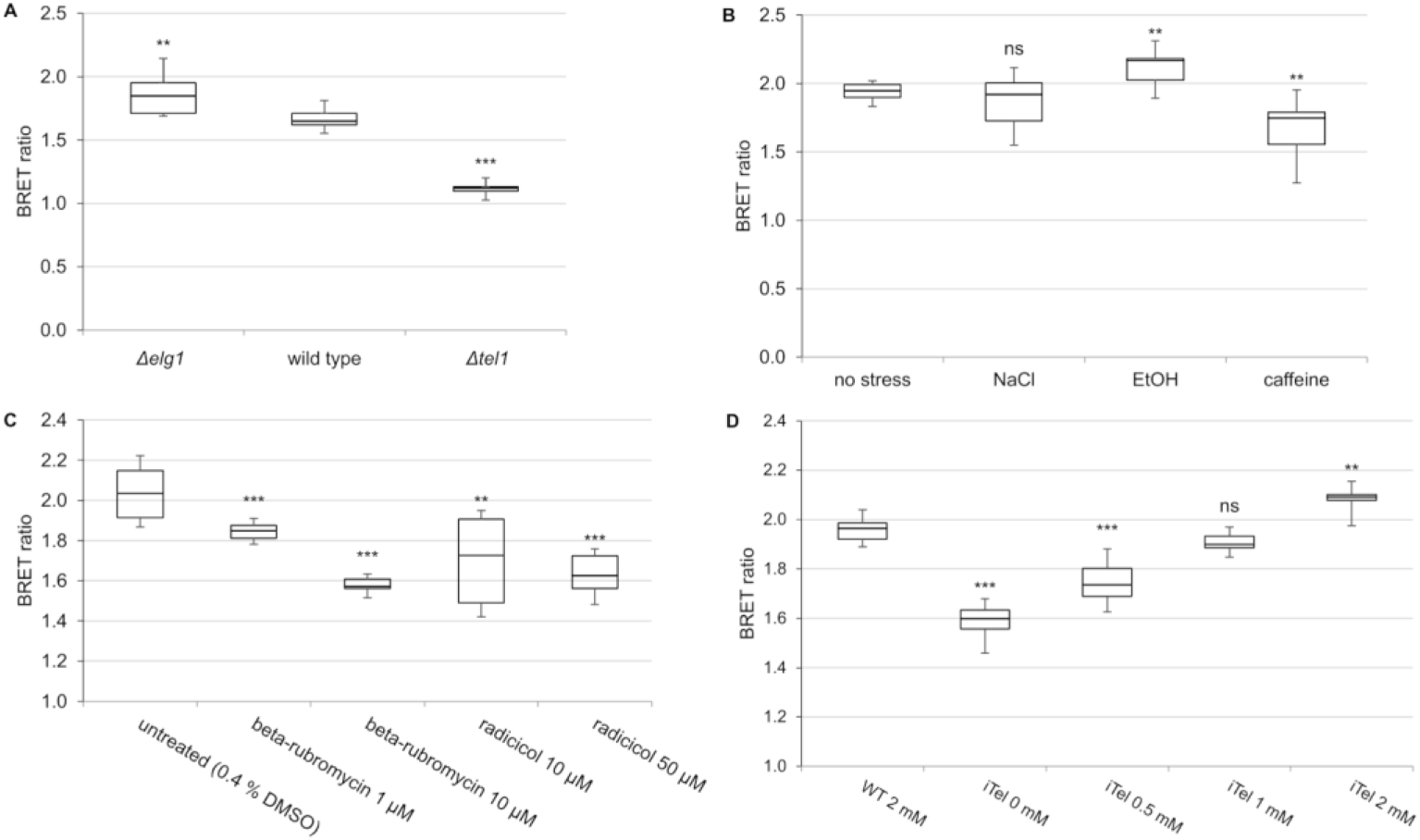
BRET ratios under different telomere length regulation conditions. (A) Deletion strains *Δelg1*, *Δtel1* and wild type (three biological replicates, each with three technical replicates). (B) Wild type strain cultured under stress conditions: Without additions (no stress), in the presence of 0.5 M NaCl, 5 % (v/v) EtOH and 10 mM caffeine (six technical replicates). (C) Wild type strain cultured in presence telomerase inhibitors: 0.4 % DMSO (untreated) or two different concentrations of the beta-rubromycin and radicicol (two biological and six technical replicates). (D) Inducible *EST2* (iTel) strains in the presence of varying concentrations of cyanamide as inducer in comparison to wild type (WT) in the presence of the maximum concentration of cyanamide (six technical replicates). ns - no significant difference, ** *p*≤0.01, *** *p*≤0.001 vs. wild type/untreated (one-sided Student’s *t*-test)

Thus, the BRET ratio in our system qualitatively followed the telomere length variations of the deletion strains reported before^21^.

Next, we aimed to recapitulate the effects of stress factors previously shown to influence the telomere length in *S. cerevisiae*: Studies have shown that 5 % ethanol (EtOH) induces telomere elongation, and 10 mM caffeine leads to telomere shortening^22^. Osmotic stress induced by 0.5 M sodium chloride (NaCl) served as a control as it is not expected to affect telomere length significantly.

As shown in Fig 1B, the BRET ratio was not significantly different in the cells exposed to 0.5 M NaCl, but was significantly higher in cells grown in the presence of 5 % EtOH and significantly lower in cells grown in the presence of 10 mM caffeine, compared to untreated cells grown in rich medium (yeast peptone dextrose - YPD 1.94 ± 0.04, NaCl 1.87 ± 0.14, EtOH 2.12 ± 0.10, caffeine 1.67 ± 0.15, 95 % CI).

Next, we tested the effect of two reported telomerase inhibitors. Beta-rubromycin, a broad reverse transcriptase inhibitor, also inhibits yeast and human telomerase^23^. The inhibiting effect of radicicol on telomerase was discovered in a yeast based screen^19^. Fig 1C demonstrates that our system detected a dose dependent, significant decrease in BRET ratio in the presence of both inhibitors (YPD with 0.4 % DMSO as control 2.04 ± 0.09, 1 µM beta-rubromycin 1.85 ± 0.03, 10 µM beta-rubromycin 1.58 ± 0.03, 10 µM radicicol 1.70 ± 0.15 and 50 µM radicicol 1.63 ± 0.07).

In summary, our results are in line with the previously shown effects of stress factors^22^ and telomerase inhibitors^19,23^ on telomere length, demonstrating the system’s ability to qualitatively detect differences in telomere length and its feasibility for compound screening.

Yeast tightly regulates its telomerase system and keeps its telomeres at a stable length under normal conditions^24^. That makes it a useful model for embryonic, stem or cancer cells^25^. To mimic telomere shortening in human somatic cells during successive divisions, we engineered an inducible telomerase strain. The native promoter of the telomerase reverse transcriptase *EST2* gene^26^ was replaced by the cyanamide-inducible promoter of the *DDI2* gene^27^. The strain is referred to hereafter as iTel (inducible telomerase).

Fig 1D illustrates the dose dependent induction of *EST2* with different cyanamide concentrations. When telomerase was absent or low (0 and 0.5 mM cyanamide), the iTel strain showed significantly lower BRET ratios compared to the wild type strain treated with the maximum induction concentration of 2 mM cyanamide. At 1 mM cyanamide, no significant difference between the wild type strain and induced iTel strain was observed. Notably, strong induction at 2 mM cyanamide led to significantly higher BRET ratios compared to the wild type. The results indicate that the assay detected both telomere erosion in the absence of telomerase and telomere elongation, when telomerase was induced.

To quantify the correlation between telomere length and BRET ratio, telomere length was measured by long-read sequencing as a reference method. The previously published algorithm determines the length for each chromosome end separately and counts telomeres that can be mapped to specific subtelomeres of a reference genome^16^. Since BRET ratios were expected to correlate with average total telomeric repeats per cell, an algorithm was developed that identifies and measures all uninterrupted telomeric repeat streaks in sequencing data and calculates average telomere length and variation. The algorithm is explained in detail in the methods section.

Cells from the *Δelg1*, *Δtel1*, iTel (uninduced and induced with 2 mM cyanamide) and wild type strains (grown YPD without additions, with 0.5 M NaCl or 10 mM caffeine) were harvested directly after BRET measurements, and the resulting BRET ratios were correlated with the average telomere lengths.

Fig 2A illustrates the linear correlation between telomere length and BRET ratio. The true correlation coefficient between the measured variables was estimated 0.96 (95 % CI: 0.90 – 0.99) using parametric bootstrapping.

**Fig 2.**
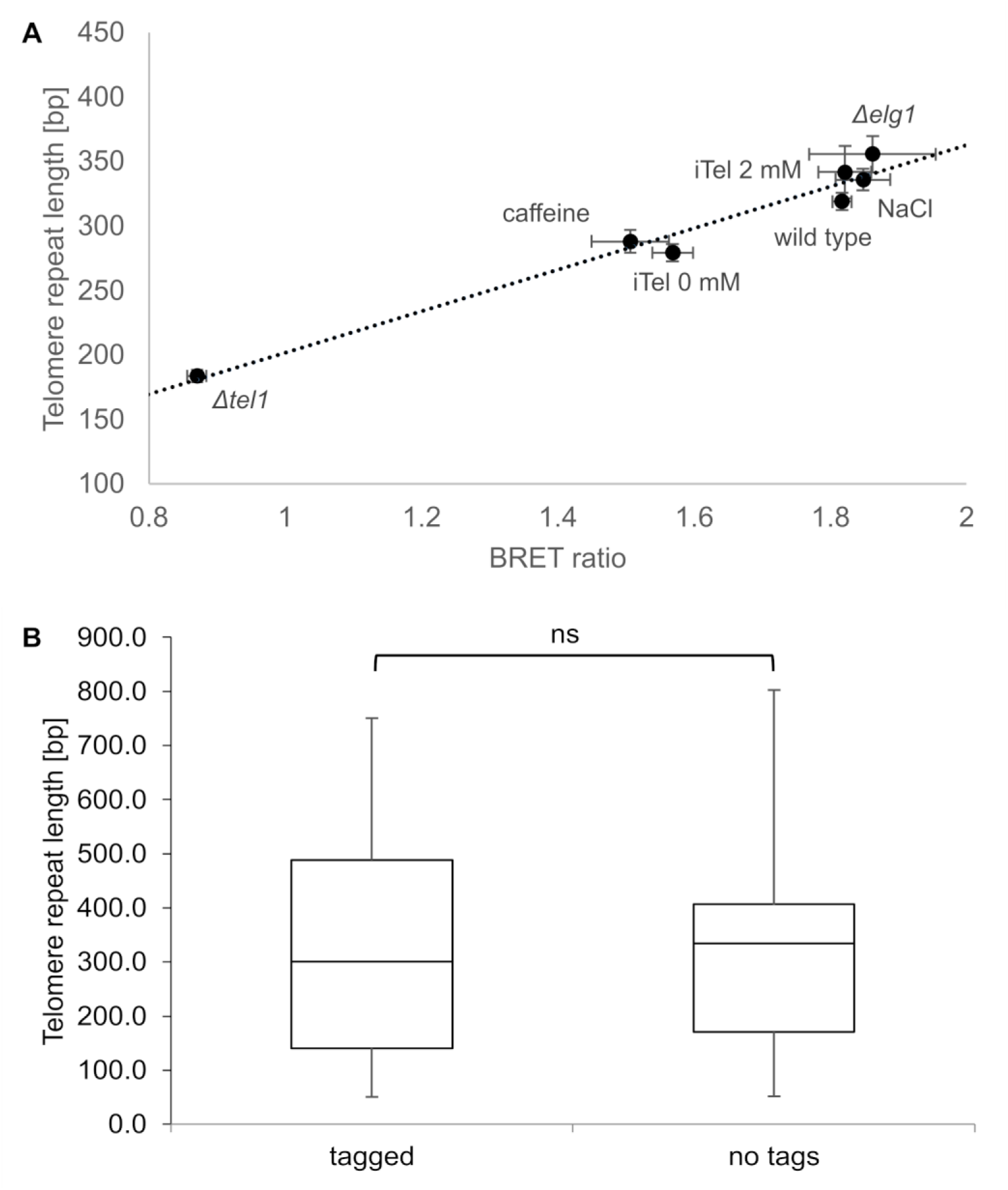
Correlation of BRET ratios and telomere length measured by nanopore sequencing. (A) BRET ratios (8 technical replicates each) of *Δelg1*, *Δtel1*, iTel (uninduced and induced with 2 mM cyanamide) and wild type strains (grown in YPD without additions, with 0.5 M NaCl and 10 mM caffeine) plotted against telomere length determined by nanopore sequencing. Error bars represent standard error of mean (SEM). (B) Telomere length of tagged and untagged *S. cerevisiae* strains after 9 d cultivation in YPD determined by nanopore sequencing.

After 9 days (∼40 generations) long-read sequencing of untagged and tagged wild type cells was performed. The sequencing results showed no significant difference between the strains (Fig 2B). This shows that the tagging of Rif2 and Rap1 did not influence equilibrium telomere length.

Previous studies using terminal restriction fragment (TRF) analysis demonstrated that *Δelg1* possess longer telomeres and *Δtel1* shorter telomeres compared to wild type^21^. Our data supports these findings and suggest that the difference in telomere length between wild type and *Δelg1* is smaller than between wild type and *Δtel1* telomeres (Fig 1A). This was confirmed by the sequencing data (Table 1).

**Table 1:** BRET ratio and corresponding telomere lengths (determined by nanopore sequencing) of tagged *S. cerevisiae* strains presented with 95 % CI.

| Tagged strain | Telomere length | BRET ratio |
| --- | --- | --- |
| *Δelg1* | 355 ± 28 bp | 1.86 ± 0.18 |
| wild type | 318 ± 13 bp | 1.82 ± 0.01 |
| *Δtel1* | 183 ± 9 bp | 0.87 ± 0.03 |
| iTel 0 mM | 279 ± 13 bp | 1.57 ± 0.06 |
| iTel 2 mM | 341 ± 40 bp | 1.82 ± 0.08 |
| NaCl | 335 ± 16 bp | 1.85 ± 0.08 |
| Caffeine | 287 ± 17 bp | 1.51 ± 0.06 |

The observed correlation between telomere length and BRET ratio provides evidence that the system enables quantitative assessment of telomere length.

Linear regression analysis of BRET ratio and telomere length revealed a direct proportionality, enabling estimation of telomere length from BRET ratio within the tested range of 183 to 355 bp (Equation 2). Notably, Equation 2 is experiment-specific and must be determined for each specific experimental setup.

To characterize the system in more detail, we evaluated the influence of cell density and incubation times on the BRET ratio. Although both variables markedly affected the absolute luminescence intensity (data not shown) they had no significant influence on the BRET ratio. BRET ratios were measured across a more than 100-fold range of cell densities (OD_600_ 0.02 to 3) using serial dilutions of *Δelg1*, wild type, *Δtel1* and a negative control. Furthermore, BRET ratios were monitored over a time course from 6 to 70 minutes after addition of the luminescence substrate. A wild type strain expressing the Rluc-Rif2 in the absence of a fluorescence acceptor was used as a negative control.

Although luminescence intensity is influenced by cell density and incubation time of the luminescence substrate (data not shown), the BRET ratio remained stable for each strain demonstrating that the system can measure telomere length independently of these factors (Fig 3A and B).

**Fig 3.**
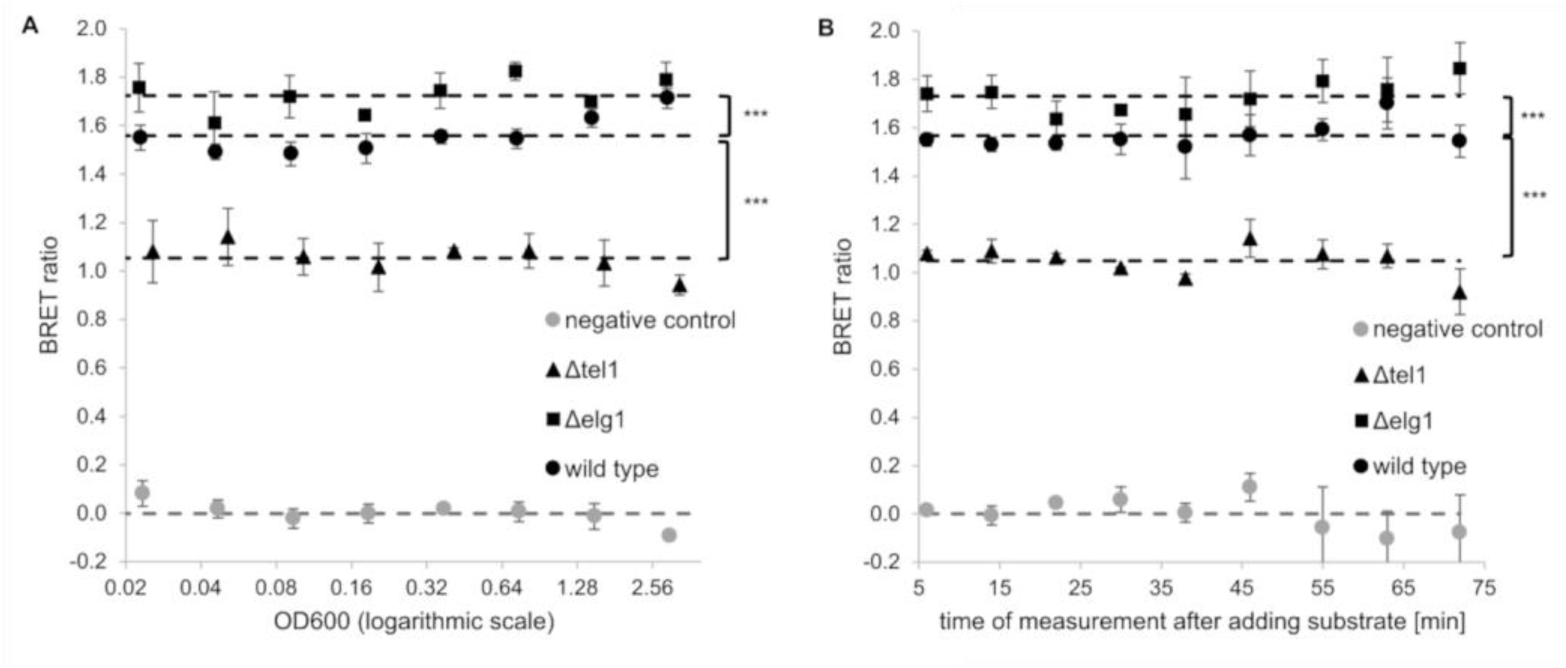
Variation of BRET ratio with cell density and time of measurement. (A) Deletion strains *Δelg1*, *Δtel1,* wild type and negative control in relation to OD_600_ (3 technical replicates). (B) *Δelg1*, *Δtel1,* wild type and negative control measured continuously and plotted against time (in minutes) following injection of the luminescence substrate. *** *p*≤0.001 vs. wild type (one-sided Student’s *t*-test), mean value depicted as dashed line

## DISCUSSION

Methods to measure telomere length, such as terminal restriction fragment (TRF) analysis^13^, fluorescence in-situ hybridization (FISH)^14^, qPCR^15^ and long-read sequencing^16^ are time, labor and/or cost intensive. Alternatively, several methods assess telomerase activity rather than directly measuring telomere length, such as the telomeric repeat amplification protocol (TRAP)^28^. While TRAP assays enable fluorescence or luminescence-based readouts, they are not high-throughput capable^29^ and, importantly, do not assess telomere length, limiting their ability to detect telomerase-independent effects.

The system presented in this study is able to quickly and efficiently measure telomere length in living *S. cerevisiae* cells. It is based on BRET between the two tagged telomeric proteins Rap1 and Rif2. Our hypothesis that the ratio of bound to unbound Rap1/Rif2 reflects the abundance of Rap1 binding sites (i.e. telomeric repeats) and, thus telomere length, was supported by several experimental findings:

To determine whether the BRET ratio correlates with telomere length, we measured the BRET ratio in genomically tagged *S. cerevisiae* wild type, *Δelg1* and *Δtel1* deletion strains^32^. Previous TRF analyses have shown that the deletion of *ELG1* leads to elongated telomeres, whereas the deletion of *TEL1* results in shorter telomeres than in wild type^21^. Furthermore, we exposed the strains to stress factors, previously reported to disrupt telomere length homeostasis^22^ and compounds reported to inhibit telomerase^19,23^. Our system detected qualitative differences in telomere length that were in line with the reported effects.

To determine whether BRET ratios also correlate quantitatively with average telomere length we developed an algorithm that identifies and measures telomeric repeats in long-read sequences.

Using this algorithm on nanopore sequencing data, a linear correlation between BRET ratio and average telomere length could be determined. This demonstrated that the system measures telomere length quantitatively. The experiment also confirmed our observation (Fig. 1A) that the difference in telomere length between *Δelg1* and wild type is smaller than that between *Δtel1* and wild type.

It has been shown that Rif1 and Rif2 form a larger telomeric architecture by connecting multiple telomere bound Rap1 molecules. These multivalent Rap1 interactions are necessary for correct telomere length homeostasis *in vivo* and lead to a preferred Rap1 binding to arrays of more than three over single Rap1 binding sites *in vitro*^20^. The clear correlation of Rap1/Rif2-binding and telomere length, demonstrated in this study, shows Rap1/Rif2-binding is facilitated or stabilized on arrays of Rap1 binding sites. This finding could be transferable to other multivalent DNA-protein complexes like the DNA damage response.

Since the interaction of Rap1 with Rif2 (and Rif1) is necessary for telomere length homeostasis and the disruption of the Rap1-Rif2 binding modules leads to telomere elongation^20^, it was necessary to determine whether the tagging of Rap1 and Rif2 close to the binding domains of these proteins influenced the equilibrium telomere length of the yeast strains. We determined telomere length in untagged and tagged yeast cells by nanopore sequencing. There was no statistically significant difference in the length distribution of the sequenced telomeres. This allows the conclusion that the used tags have no significant influence on the binding between Rap1 and Rif2 and thus telomeric architecture and length regulation.

In general, the BRET ratios showed a higher variability between biological replicates than between their respective technical replicates. This indicates that clonal populations of yeast cells develop different average telomere lengths after less than ten generations under the same cultivation conditions. This variation in telomere length between clonal cultures was more pronounced in the *Δelg1* mutant than in wild type or *Δtel1* cells (see Fig 1A). This could be explained by the observation that *Δelg1* leads to increased genomic instability^31^.

The highest variability between clones was detected in the iTel mutants, both in the presence and absence of telomerase induction. We suggest that this variability may reflect a disruption of the tightly regulated telomere length homeostasis due to non-physiological regulation and levels of telomerase reverse transcriptase expression.

The iTel experiments demonstrated that telomere length was reduced compared to wild type when telomerase was not induced or induced at low cyanamide concentrations (0 and 0.5 mM cyanamide). Telomere length recovered to wild type levels at 1 mM cyanamide. Notably, strong induction of the *EST2* gene with 2 mM cyanamide for seven days (corresponding to more than 30 generations) resulted in significantly longer telomeres than those observed in wild type cells.

These findings suggest that overexpression of telomerase reverse transcriptase gene *EST2* can disturb the equilibrium between telomere shortening and lengthening. In wild type cells this balance is primarily maintained by preferential recruitment of the telomerase holoenzyme to critically short telomeres^18^. Both *TLC1* (telomerase RNA component) and *EST2* are naturally expressed at low levels, particularly in haploid *S. cerevisiae* cells^32^. This limited availability might serve as an additional regulatory mechanism, ensuring that telomere length remains within a defined range. Thus, overexpression of *EST2* could shift the equilibrium toward elongation, resulting in telomere lengthening beyond wild type levels over extended cultivation.

So called survivor cells in yeast cultures that continue to proliferate after prolonged cultivation in the absence of telomerase rely on alternative lengthening of telomeres (ALT)^33^. A similar phenomenon is known from a subgroup of human cancers (especially tumors of mesenchymal origin) that normally have a poor prognosis for patients^34^. The iTel system could be a valuable model to screen for telomere shortening or destabilizing compounds effective against these cancers. Additionally, using inducible telomerase, the BRET system opens possibilities to screen for compounds that decelerate the erosion of telomeres. These could be interesting candidates for the treatment of age-related diseases in somatic human cells.

While this system is based on *S. cerevisiae* and thus does not fully reflect the complexities of telomere biology in humans, it should be noted that both telomerase inhibitors used in this study are effective in yeast and humans. Moreover, the system allows the detection of mechanisms of stabilizing or destabilizing telomeric DNA independent of telomerase or other enzymes. We expect these kinds of mechanisms to have a higher potential of being transferable to human cells than specific activators or inhibitors of enzymes.

The main advantage of using BRET, a ratio of two luminescence emission intensities, is its independence from cell density. This property increases its reliability for compound screening since cell density is affected by compound effects, statistical variability, and differences in cell inoculum concentration. The ability of measuring in a wide range of OD_600_, demonstrated in Fig 3A, saves the steps of measuring and equalizing the density in those experiments.

Furthermore, BRET ratios remained stable for over one hour of measurement (Fig. 3B), allowing the study of telomere dynamics over time in living cells.

In summary, we developed a novel method to quantify telomere length in living cells. This fast and cost-efficient system enables further research on telomere regulation and the screening of large-scale compound libraries.

## MATERIALS AND METHODS

### RESOURCES TABLE

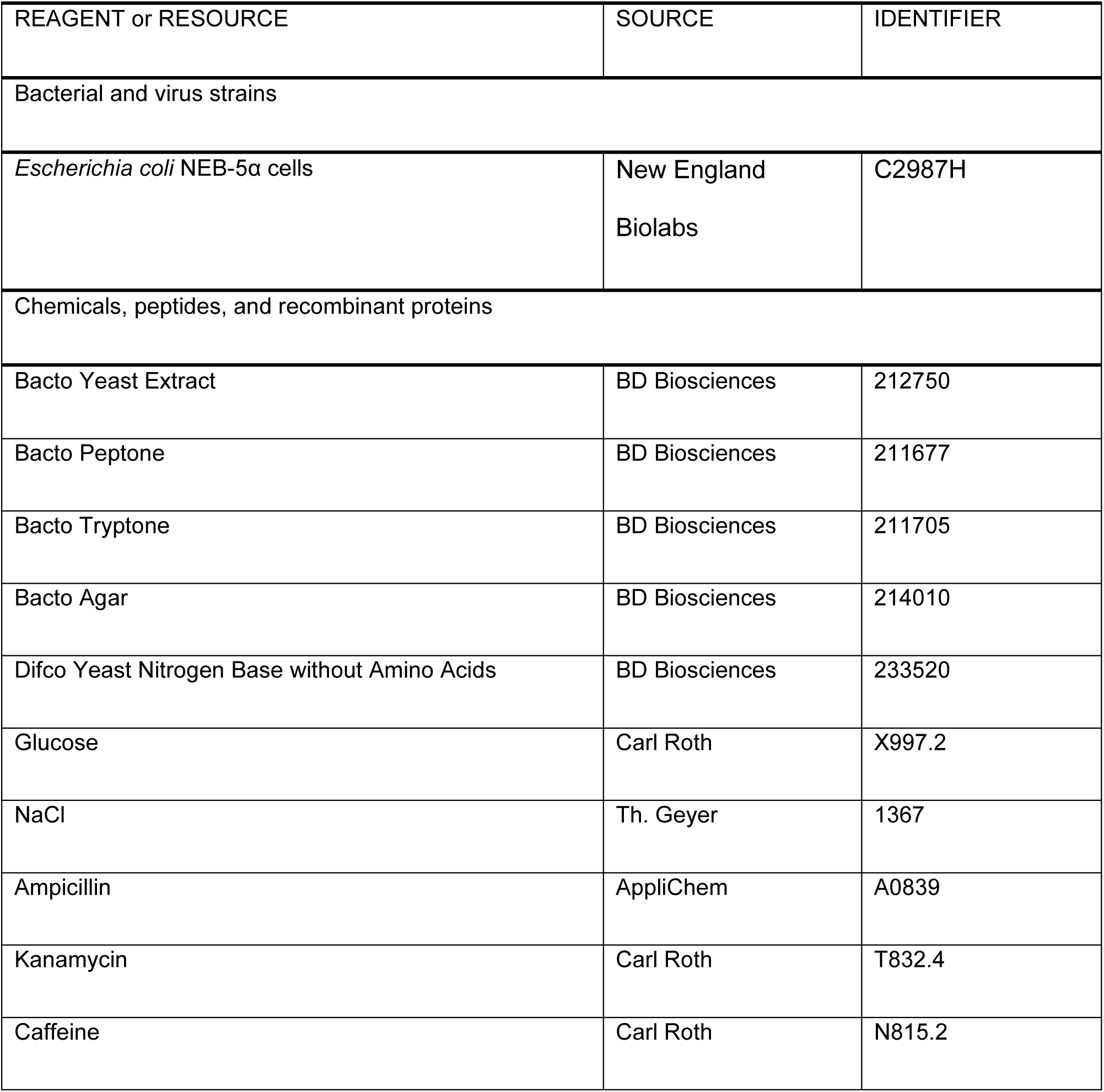
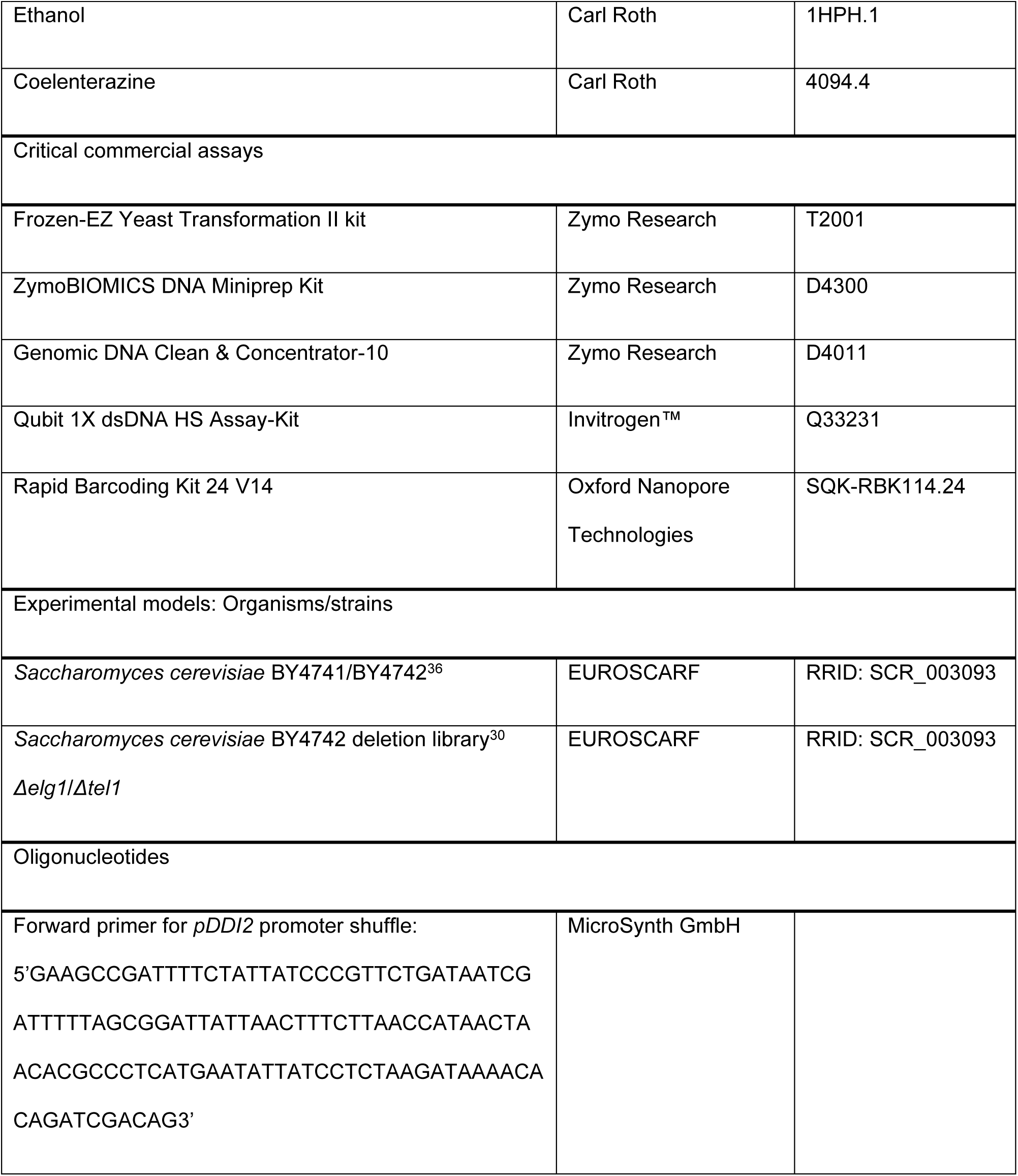
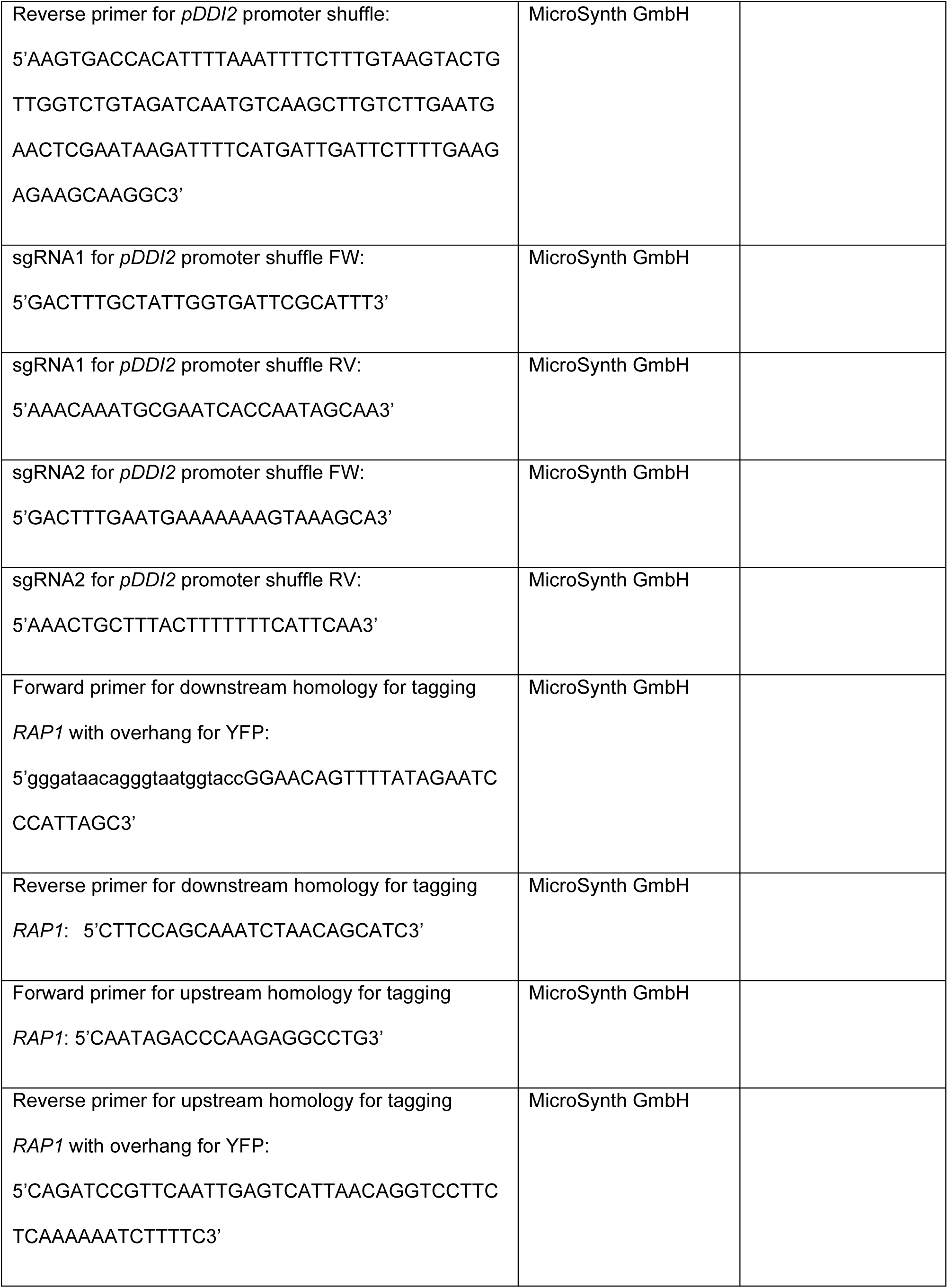
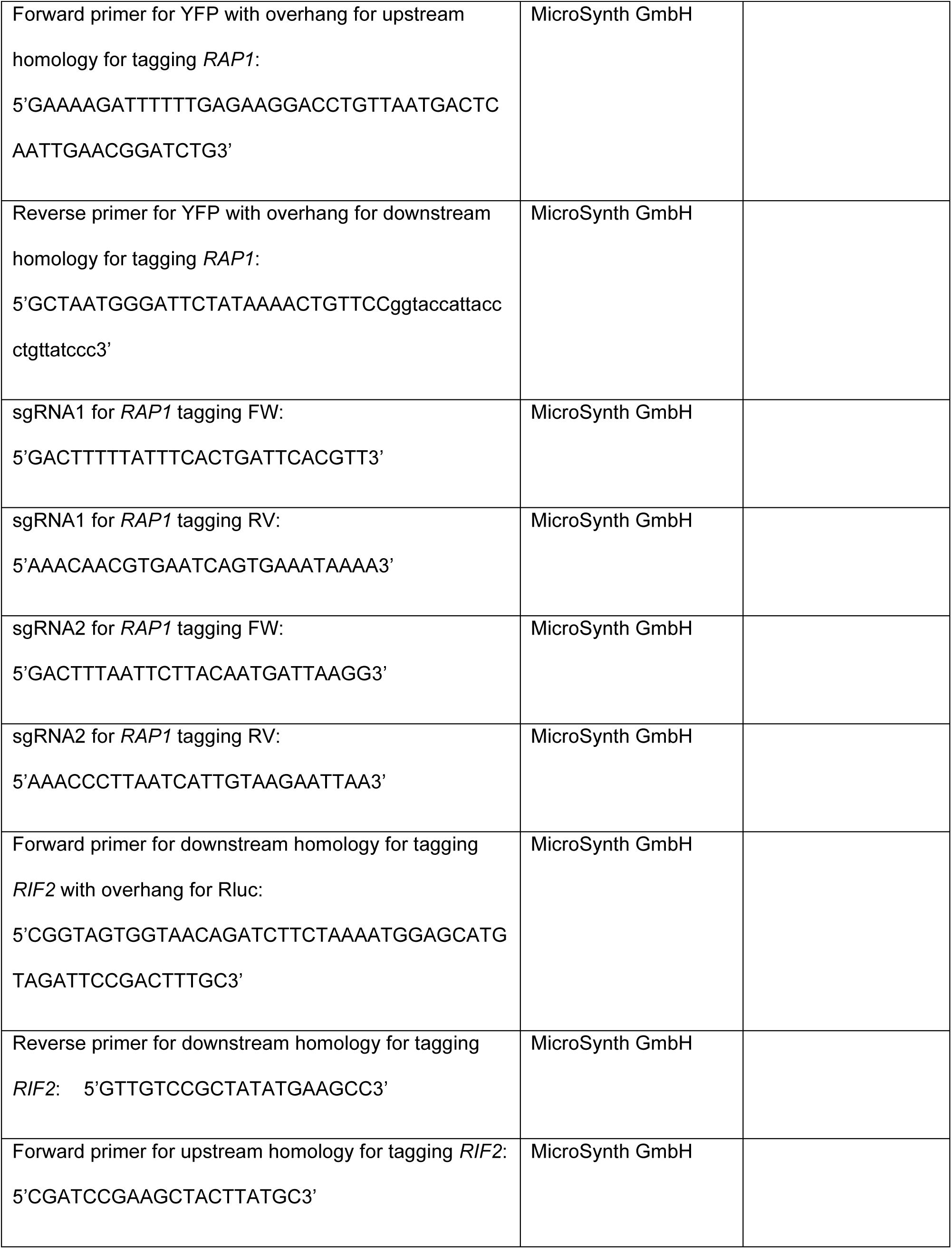
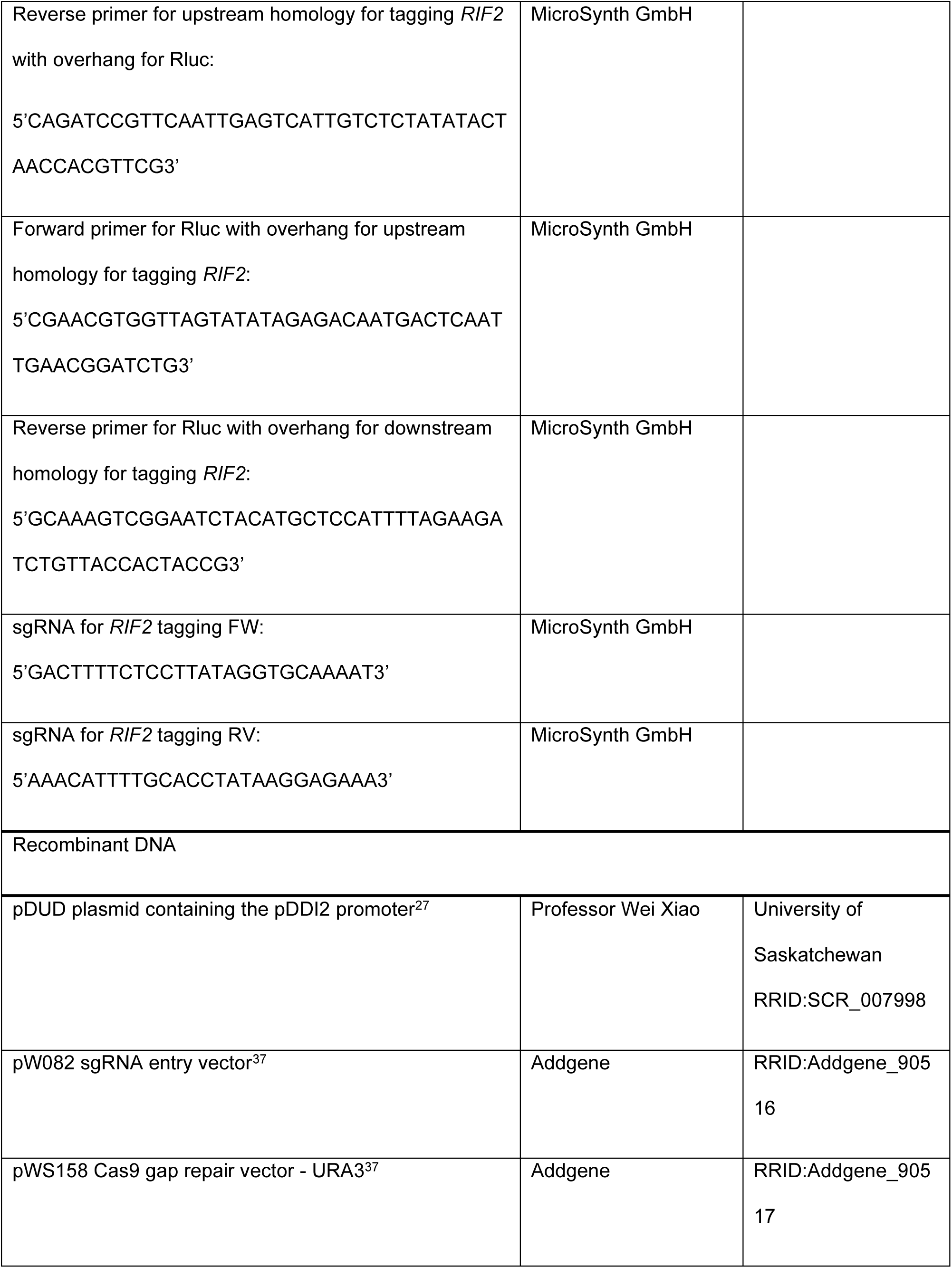
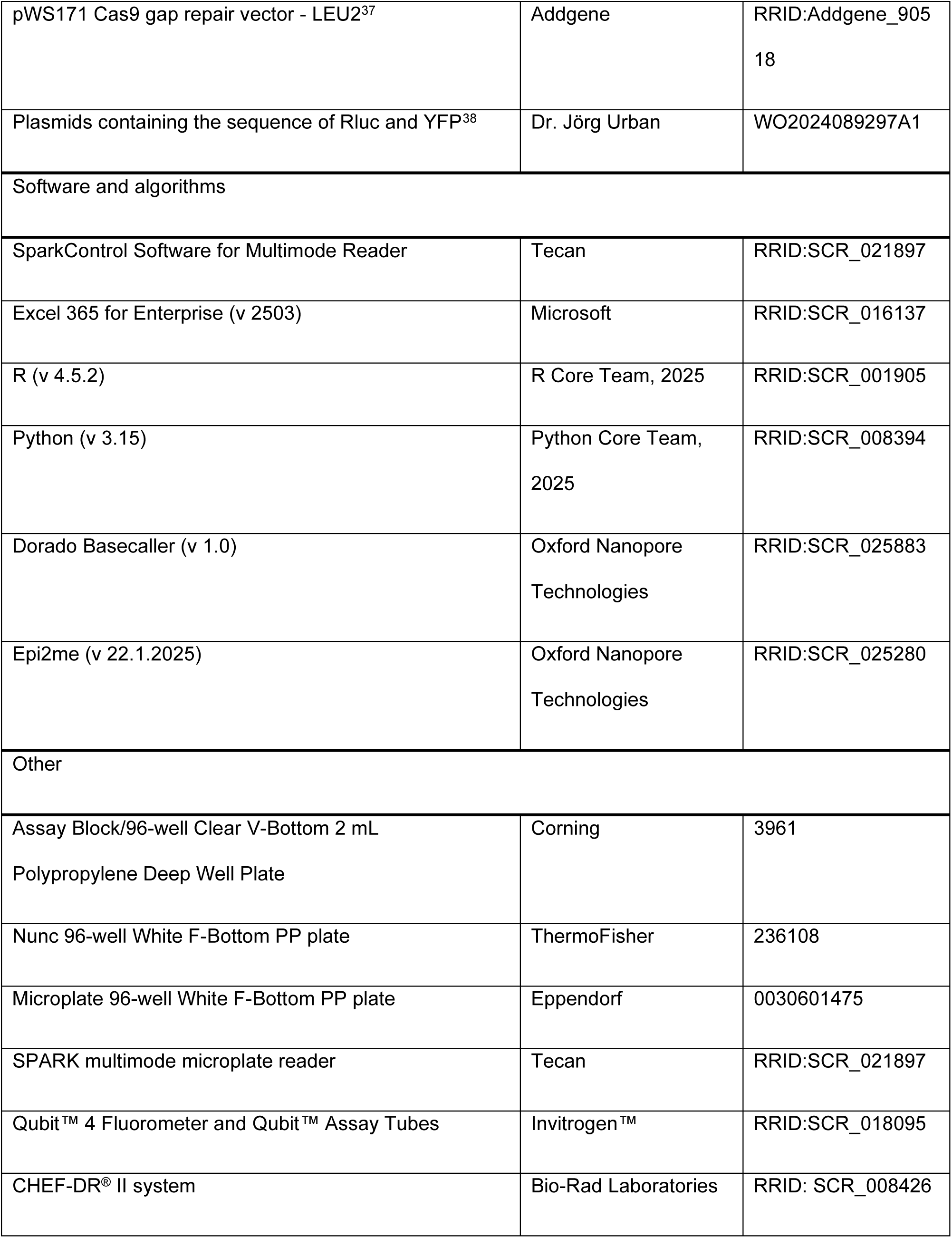
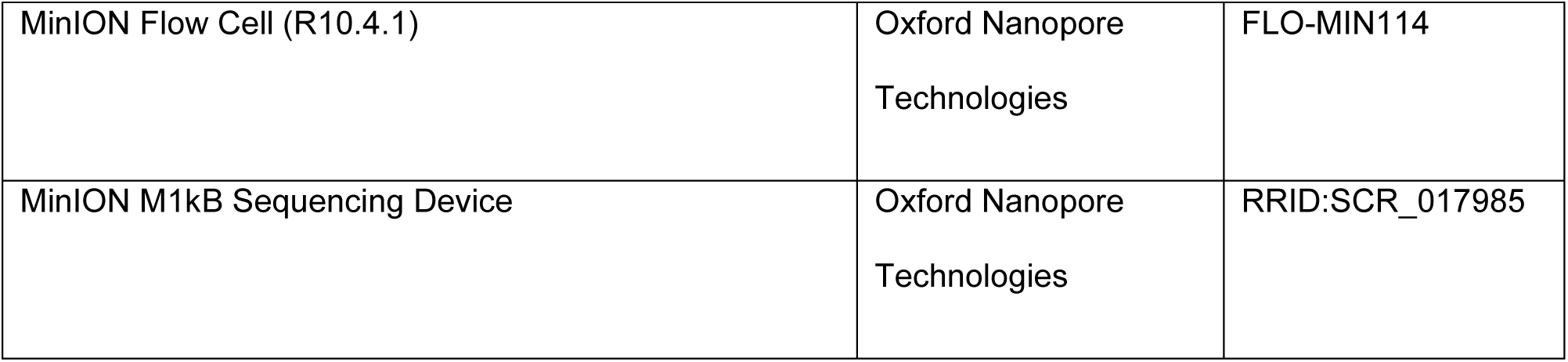

### Genomic modification of yeast

All modifications were carried out by CRISPR/Cas9 assisted, marker-free integration into the *S. cerevisiae* BY4741^36^ strain (wild type) or the deletion strains^30^ *Δelg1* (shorter telomeres) and *Δtel1* (longer telomeres)^22^. The MoClo-Toolkit vector system^37^ was used to introduce two sgRNAs targeting each genomic site to be modified. Overlap extension PCR was employed to create linear insertion constructs.^39^. The cassettes contained the insert flanked by 300 to 600 bp of homology to the target region, including silent mutations that altered the sgRNA target sequence.

The native *EST2* promoter was replaced with the inducible *DDI2* promoter in the same manner. The nucleotide sequence encoding *Renilla reniformis* luciferase (Rluc, BRET donor) was fused to the N-terminus of *RIF2*, and the sequence of a modified Yellow Fluorescent Protein (YFP, BRET acceptor) was added to the C-terminus of *RAP1*. This design enables a transient interaction with Rluc, increasing BRET efficiency^38^.

The linearized Cas9- and two sgRNA-entry vectors were co-transformed with the PCR repair DNA construct using the LiAc-method^40^ or the Frozen-EZ Yeast Transformation II kit (Zymo Research).

These modifications resulted in tagged wild type *S. cerevisiae*, tagged *Δelg1* and tagged *Δtel1* strains used in our system.

### Culture conditions and preparation for measurement

Yeast was streaked from glycerol freeze stocks on YPD agar (5 % yeast extract, 10 % peptone, 2 % glucose, 2 % agar, pH 5), single colonies were transferred to YPD medium w/o supplementation and cultured in an orbital shaking incubator at 30 °C and 230 rpm in 50 ml Erlmeyer flasks (for harvest and DNA-extraction) or at 800 rpm in deep-well plates.

Cells were passaged by 1:1000 dilution in fresh YPD medium every 2 to 3 days. Cultivation times are indicated in the text.

### Measurement of BRET ratios

Stationary cultures (after 2 to 3 days of culturing) were diluted 1:4 in YPD medium and dispensed 200 µl per well for each technical replicate (see Results) into white 96-well plates (ThermoFisher, USA) for measurement.

At least 15 min before measurement coelenterazine (2.5 mM in EtOH) was diluted 1:100 in YPD medium. Of this diluted substrate 50 µl per well (final conc. 5 µM) were added and mixed by pipetting up and down directly before measurement.

The plate reader was preheated to 30 °C. Cells were incubated for 5 min by shaking (200 RPM) with the coelenterazine before measurement. Integration time was 1 s per wavelength.

BRET ratio is calculated as emission intensity in the wavelength band 505 to 590 nm (YFP) divided by emission intensity in the wavelength band 430 to 485 nm (Rluc) minus BRET ratio of negative control (luciferase only).

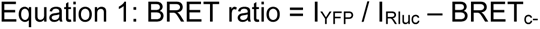

where:

I_YFP_ - photon count 505 to 590 nm,

I_Rluc_ - photon count 430 to 485 nm,

BRET_c-_ - BRET ratio of a control strain with only the donor construct (Rluc-RIF2) and no acceptor measured in the same experiment.

### Statistics

The confidence intervals, *p*-values (using one-sided Student’s *t*-test) were calculated in Excel 365 for Enterprise (v2503). If not mentioned otherwise, *t*-tests were two independent samples (unpaired) and one-sided where a clear direction of the treatment effect was expected. Normal distribution of the values was assumed, and the variance of the populations was equal.

The linear regression and estimation of the Pearson’s correlation coefficient in Fig 2 was carried out using R.

To account for variance in both variables, correlation between BRET ratio and telomere length was estimated using a parametric bootstrap approach. For each data point, 10,000 synthetic datasets were generated by sampling from normal distributions centered at the measured mean and scaled by the observed standard error of the mean (SEM). For each bootstrap iteration, Pearson’s correlation coefficient was computed, and the distribution of coefficients was used to determine the mean correlation and 95% confidence interval.

The linear regression yielded the following equitation for this experiment:

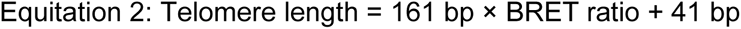

### Nanopore sequencing

Telomere tagging and nanopore sequencing were carried out as previously described^16^, with the following changes: Commercial kits were used for DNA extraction and purification according to the manufacturer’s instructions (Zymo Research, D4300 and D4011). For tagged DNA samples, the Rapid Barcoding Kit (SQK-RBK114.24) was used. The sequencing run was performed on a MinION flow cell (R10.4.1) for 40 h using MinKNOW software (2v 4.06.16) resulting in 13- to 33-fold coverage per barcode.

### Determining telomere length from long-read sequences

The algorithm that calculates average telomere length was written in Python (v 3.15). It scans concatenated (seqkit concat) fastq files for telomeric sequences that generally have the form C_1-3_A / TG_1-3_. The algorithm excludes pure TG/AC repeats as well as incomplete telomeric reads, where the sequence ends within the telomeric repeats. It stops counting when more than two A or C (on the TG rich strand) / T or G (on the AC rich strand) within a 15 bp window or more than 3 T or 5 G (A/C on the opposite strand) in a row are found. It generates a list with an index, the length and the sequence +/- 50 bp of each detected telomere. Based on this list statistical analysis can be carried out.

The python code is available on Github and can be used freely (with attribution): https://github.com/Felix-FrankeLab/Telofinder.git

## ACKNOWLEDGMENTS

We thank Professor Wei Xiao, Department of Biochemistry, Microbiology and Immunology, University of Saskatchewan, Canada for providing the plasmid with the DDI2 promoter.

We thank Peter Lachmann for helpful support of sequencing data analysis and critical reading of the manuscript.

## AUTHOR CONTRIBUTIONS

F.R. Conceptualization, Data curation, Formal analysis, Funding acquisition, Investigation, Methodology, Resources, Validation, Visualization, Writing – original draft

H.M.R. (nanopore sequencing) Investigation, Writing – review & editing

J.U. (BRET Tags) Methodology, Resources, Writing – review & editing

J.F. Investigation, Methodology, Funding acquisition, Project administration, Resources, Supervision, Writing – review & editing

